# Nearly all new protein-coding predictions in the CHESS database are not protein-coding

**DOI:** 10.1101/360602

**Authors:** Irwin Jungreis, Michael L. Tress, Jonathan Mudge, Cristina Sisu, Toby Hunt, Rory Johnson, Barbara Uszczynska-Ratajczak, Julien Lagarde, James Wright, Paul Muir, Mark Gerstein, Roderic Guigo, Manolis Kellis, Adam Frankish, Paul Flicek, The GENCODE Consortium

## Abstract

In a 2018 paper posted to bioRxiv, Pertea et al. presented the CHESS database, a new catalog of human gene annotations that includes 1,178 new protein-coding predictions. These are based on evidence of transcription in human tissues and homology to earlier annotations in human and other mammals. Here, we reanalyze the evidence used by CHESS, and find that nearly all protein-coding predictions are false positives. We find that 86% overlap transposons marked by RepeatMasker that are known to frequently result in false positive protein-coding predictions. More than half are homologous to only nine *Alu*-derived primate sequences corresponding to an erroneous and previously withdrawn Pfam protein domain. The entire set shows poor evolutionary conservation and PhyloCSF protein-coding evolutionary signatures indistinguishable from noncoding RNAs, indicating lack of protein-coding constraint. Only four predictions are supported by mass spectrometry evidence, and even those matches are inconclusive. Overall, the new protein-coding predictions are unsupported by any credible experimental or evolutionary evidence of function, result primarily from homology to genes incorrectly classified as protein-coding, and are unlikely to encode functional proteins.

## Introduction

The complete and accurate representation of all the genes in the human genome is critically important for both the investigation of genome biology and the medical interpretation of genetic variation. Errors in protein-coding gene annotations can result in false predictions of mutations with severe phenotypic consequences in a clinical setting (MacArthur et al. 2012).

Consequently, a reference gene catalog must only represent a given locus as protein-coding when that is believed to be the most likely interpretation of its functionality based on all the available evidence.

Identification of all transcribed loci and the determination of which of those loci are translated into functional proteins is a core remit of large-scale gene annotation projects such as Ensembl/GENCODE (henceforth GENCODE) and RefSeq, which began producing reference annotation on the human genome sequence approximately 15 years ago (Harrow et al. 2012; Zerbino et al. 2018; O’Leary et al. 2016). For GENCODE, the distinction between *bona fide* protein-coding genes and other gene types involves the expert curation of all loci on a case-by-case basis, incorporating computational pipelines and experimental data alongside targeted experimental validation procedures. Projects like GENCODE also actively follow advances in genome annotation because they recognize the critical need to achieve the highest possible accuracy for the human gene set. The GENCODE Consortium welcomes all efforts to push the state of the art and has worked closely with and evaluated the outcomes of numerous previous and ongoing studies.

Expert manual curation has resulted in high-confidence annotations for approximately 20,000 protein-coding genes in GENCODE, RefSeq, and UniProtKB/Swiss-Prot (The UniProt Consortium 2017). These databases do not claim to have a “final” count of protein-coding genes, and each remains a work in progress, in fact disagreeing from each other on the protein-coding status of approximately 2,700 loci (Abascal et al. 2018).

Numerous recent efforts have sought to detect previously undiscovered (“novel”) protein-coding genes using experimental and computational tools. These include: mass spectrometry (Slavoff et al. 2013; Gascoigne et al. 2012; Kim et al. 2014; Wilhelm et al. 2014; Vanderperre et al. 2013; Mackowiak et al. 2015); ribosomal profiling (Bazzini et al. 2014; Mackowiak et al. 2015; Crappé et al. 2013); and evolutionary signatures of protein-coding constraint (Mackowiak et al. 2015; Gascoigne et al. 2012). Unfortunately, all three techniques can produce false positive annotations (Uszczynska-Ratajczak et al. 2018): lowering validation parameters in mass spectrometry analyses results in large numbers of incorrectly matched peptides (Ezkurdia et al. 2015; Nesvizhskii 2014); ribosome association is not sufficient to validate novel coding open reading frames (Verheggen et al. 2017; Guttman et al. 2013); and evolutionary signatures are dependent upon robust genomic alignments and can produce spurious signals for pseudogenes and genomic regions antisense to true coding regions (Mudge et al. In preparation).

In contrast to transcript annotations, which can still substantially improve using increasingly large RNA datasets, high-confidence annotation of functional protein-coding regions remains challenging. A key difficulty common to many diverse genome-wide searches is the needle-in-a-haystack problem, which is exemplified well by protein-coding searches. Only a small fraction of the genome is protein-coding, and the human genome has now been so well studied that any protein-coding genes waiting to be discovered are likely to be sparsely distributed and particularly well hidden. For example, they could have particularly small coding sequence, or have highly restricted expression. This means that even highly-specific methods that work well on poorly-studied genomes will have considerably more false positives than true positives when applied to the whole human genome.

With this problem in mind, the GENCODE project does not directly import predictions produced by high-throughput discovery studies into the GENCODE protein-coding gene set, but instead subjects each prospective novel coding gene to manual appraisal. This process considers the merits for coding annotation provided by the authors, but also involves the consideration of multiple orthogonal datasets alongside a potential reanalysis of the original data.

A paper recently posted on bioRxiv introduced a new database (CHESS) of human genes, identified by generating transcripts from 9,795 RNA-seq experiments from the genotype-tissue expression (GTEx) project (The GTEx Consortium 2015), and applying various filters to predict 21,306 protein-coding and 21,856 noncoding genes, as well as millions of transcripts classified as transcriptional noise (Pertea et al. 2018). The database includes 1,178 novel protein-coding gene predictions and 3,819 novel noncoding gene predictions (i.e. not already annotated in GENCODE or RefSeq). Here, we examine the 1,178 loci that were predicted to be novel protein-coding genes (henceforth “novel protein-coding predictions”, or, for brevity, “novel coding predictions”), and the criteria used to distinguish them. We find that almost all of these loci are unlikely to be protein-coding.

## Results

### CHESS criteria allow for low-quality annotations

The approach used to distinguish protein-coding from noncoding genes in the CHESS database relies on homology to previously-annotated proteins rather than evolutionary signatures or experimental evidence of translation. More specifically, transcripts are classified as protein-coding if they contain an open reading frame (ORF) of at least 60 amino acids that shows significant homology to a human or other mammalian protein-coding annotation in GenBank (Benson et al. 2013) or UniProtKB/Swiss-Prot (The UniProt Consortium 2017), that does not overlap ribosomal RNA genes or a subset of repeat elements (LINEs or LTRs), and that is at least 75% of the length of the matched protein-coding annotation. CHESS also imposes conditions on level of transcription and diverse criteria to choose among overlapping transcripts. These criteria result in 1,178 loci classified as novel protein-coding genes.

*A priori*, these criteria leave open three major sources of false positives: homology to protein-coding annotations that were themselves erroneous (thus propagating the errors of low-quality genome annotations), overlap with non-LTR and non-LINE repeat elements (e.g., SINE *Alu* repeat elements), and overlap with pseudogenes (non-functional gene remnants that bear some resemblance to their still-functional relatives).

### Little similarity to validated coding genes

Predicting protein-coding regions in a poorly-annotated genome based on homology to a high-quality “target” genome is a common technique (Zerbino et al. 2018). Such an approach is less useful for reappraising a well-annotated genome, because homologs of well-known protein-coding genes are likely to be already annotated as coding (or pseudogene).

The target protein set used for homology search in the CHESS analysis pipeline included both the expert-curated UniProtKB/Swiss-Prot protein database and the RefSeq gene catalog, which includes both manual and computationally-derived protein annotations. However, the pipeline also utilized the entire set of sequences from GenBank. It is important to note that GenBank is a repository of nucleotide sequences - a subset of which have accompanying translations - and not an expert-curated catalog of protein-coding genes. While it is possible that some predictions based on homology to lower quality annotations such as GenBank could be true protein-coding genes, it is important to note that using such annotations as the target increases the likelihood of generating false positive predictions, and these can constitute the bulk of what remains after filtering out previously annotated homologs of true coding genes. A total of 1,083 of the novel protein-coding predictions were tagged as having homology to GenBank sequences, 616 had homology to UniProtKB/SwissProt sequences and 93 to RefSeq sequences (some novel coding predictions had homology to more than one target protein).

The most striking feature of the novel protein-coding predictions is their similarity to one another. Most or the novel coding predictions appear to be homologs of other novel coding predictions (903 of them have homology to just 177 GenBank or predicted RefSeq sequences). According to the CHESS database 102 of the 1,178 novel coding predictions were classified as protein-coding based on their homology with GenBank sequence AAP34472.1, and a further 37 had homology with GenBank prediction EHH18952.1. GenBank sequence AAP34472.1 (also known as LP3428) is one member of a group of human conceptual translations derived from SVA (SINE-VNTR-*Alu*s) transposons (Hancks and Kazazian 2010) that were created in the early 2000s. SVA transposons are evolutionarily young, active, and present in about 2,700 copies in the human reference genome (Wang et al. 2005). A further 13 novel coding predictions are similar to HCA25a, another SVA transposon, and 34 are similar to one of several “FKSG” proteins annotated in GenBank, most of which are again related to SVA retroelements.

Most of the 37 novel coding predictions similar to GenBank prediction EHH18952.1 (a hypothetical protein from macaque) are also similar to the human UniProtKB/Swiss-Prot protein Q86U02 from gene *LINC00596*. *LINC00596* is annotated as noncoding in both GENCODE and RefSeq and in UniProtKB/Swiss-Prot is tagged with the lowest evidence code “unclear”. Another UniProtKB/Swiss-Prot protein (Q6UX73 from gene *C16orf89*) is also similar both to novel coding predictions and to GenBank prediction EHH18952.1. Unlike *LINC00596*, *C16orf89* is annotated as protein-coding in both GENCODE and RefSeq and has the highest UniProtKB evidence coding “protein evidence”. In fact the major isoform of *C16orf89* is well conserved in mammals and is known to be expressed in thyroid (Afink et al. 2010). However, the region of the *C16orf89* that is similar to *LINC00596* and EHH18952.1 is the product of an alternative 3’ exon. What both *LINC00596* and the alternative 3’ exon of *C16orf89* have in common is a GVQW domain.

GVQW domains were Pfam domains (Finn et al. 2014) that are now obsolete. The NCBI conserved domain database (Marchler-Bauer et al. 2017) still annotates the domain with the following description “This short domain is often found nested inside other longer domains. The function is not known, but the domain carries a highly conserved GVQW motif. It is possible that this is an AluS that has expanded in Human and Macaque genomes.” The GVQW domain was made obsolete in Pfam precisely because it is based on an *Alu* SINE element. Before it was removed, the domain was used to validate the protein-coding potential of a handful of human and other primate genes. Genes annotated with this domain are in the process of being reclassified as noncoding in the GENCODE reference set (Abascal et al. 2018), but genes *GVQW1* and *GVQW2*, both annotated with the now defunct GVQW domain, are still annotated as coding in RefSeq and UniProtKB/SwissProt.

In fact more than half of the novel protein-coding predictions in the CHESS database appear to be based on similarity to proteins with this obsolete GVQW domain: 560 are similar to just nine proteins or regions of proteins in UniProtKB/SwissProt that have a GVQW domain (**Fig. 1**, Supplementary Table 1) and 649 are either similar to one of these nine proteins, or similar to a GenBank sequence that is similar to one of the nine proteins. Thus, more than half of the novel protein-coding predictions in CHESS appear to be derived from *Alu* SINE elements.

**Figure 1.**
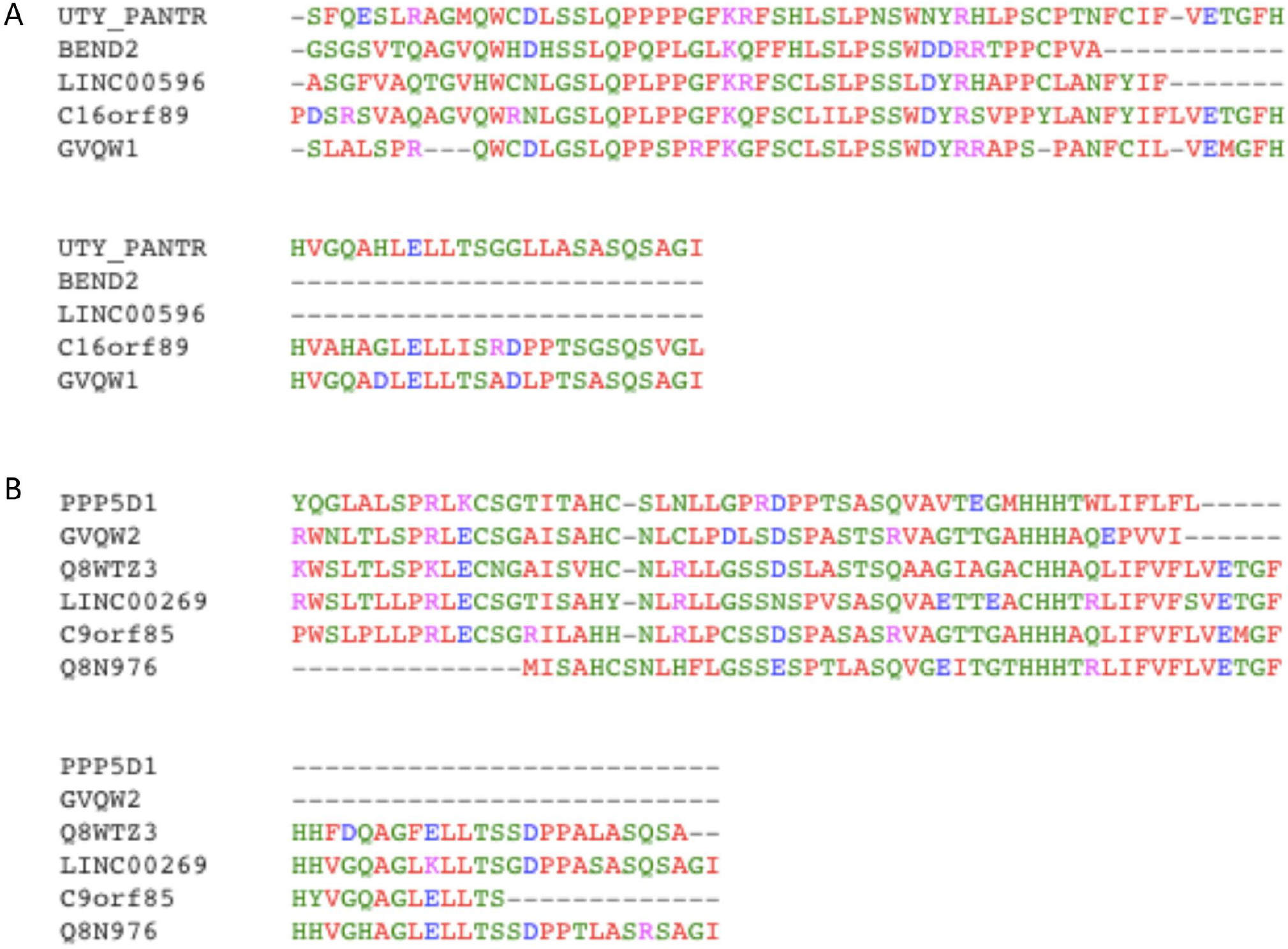
Alignment of the two types of GVQW *Alu* SINE sequences. Multiple alignments of (A) sequences similar to *GVQW1* and (B) sequences similar to *GVQW2* using KAlign (Lassmann et al. 2009). Colours indicate similarity of the amino acid. Note that the C-terminals of the transposon-derived sequences similar to *GVQW1* and those similar to *GVQW2* are homologous, but the N-terminal region is not. Identifiers that start with the letter Q are UniProt identifiers and indicate that this sequence is present in UniProt only. Four genes (*GVQW2*, *C9orf85*, *C16orf89* and *BEND2)* are annotated as coding in GENCODE. In all but the first sequence the transposon-related sequence is part of an alternative exon. *GVQW2* is already earmarked for reclassification in GENCODE.

Eight novel coding predictions are similar to the last 50 residues of UniProtKB/SwissProt protein Q96N38 (from gene *ZNF714*), which again is provided by an *Alu* element and a further eight are similar to UniProtKB/SwissProt protein Q96CB5 (*C8orf44*), a gene that was flagged with potential noncoding features (Abascal et al. 2018). The region of homology to the novel sequences precisely matches a region of *Alu* sequence in the C8orf44 coding sequence. In total *at least* 846 of the novel protein-coding predictions are similar to sequences in databases that appear to have been derived from transposons (Supplementary Table 1).

### Overlap with RepeatMasker regions

Since a majority of the novel protein-coding predictions were classified as protein-coding based on homology to transposon-associated sequences, we used RepeatMasker (Smit et al. 2013) to investigate how many of the predicted novel coding transcripts themselves overlapped interspersed repeats and low complexity DNA sequences. The ORFs of 92% of the predicted novel coding transcripts have some overlap with a RepeatMasker region, including 34% that are entirely contained in such regions. The ORFs of around 25% of GENCODE coding transcripts also overlap RepeatMasker regions, but in most cases only a small fraction of the ORF overlaps, in sharp contrast to the novel ORFs (Fig. 2). In fact, RepeatMasker predicts that 61% of all nucleotide positions in the novel ORFs are in a transposon-associated region and another 11% are in other repetitive or low complexity regions, whereas only 0.96% of all nucleotide positions in GENCODE ORFs are in a transposon-associated region and another 1.1% in other repetitive or low complexity regions. The fraction of transposon overlap with nucleotides in ORFs of APPRIS principal isoforms (Rodriguez et al. 2018), a subset of GENCODE transcripts considered most likely to represent primary functional translations based on metrics including evolutionary conservation, is just 0.37%, so most transposon overlap with GENCODE ORFs is in alternatively spliced exons that are not part of an APPRIS principal isoform. Looking more closely at the classes of repetitive regions that overlap novel protein-coding predictions we see that more than 67% overlap SINE/Alu elements and 15.6% overlap Retroposon/SVA elements (Table 1).

**Figure 2.**
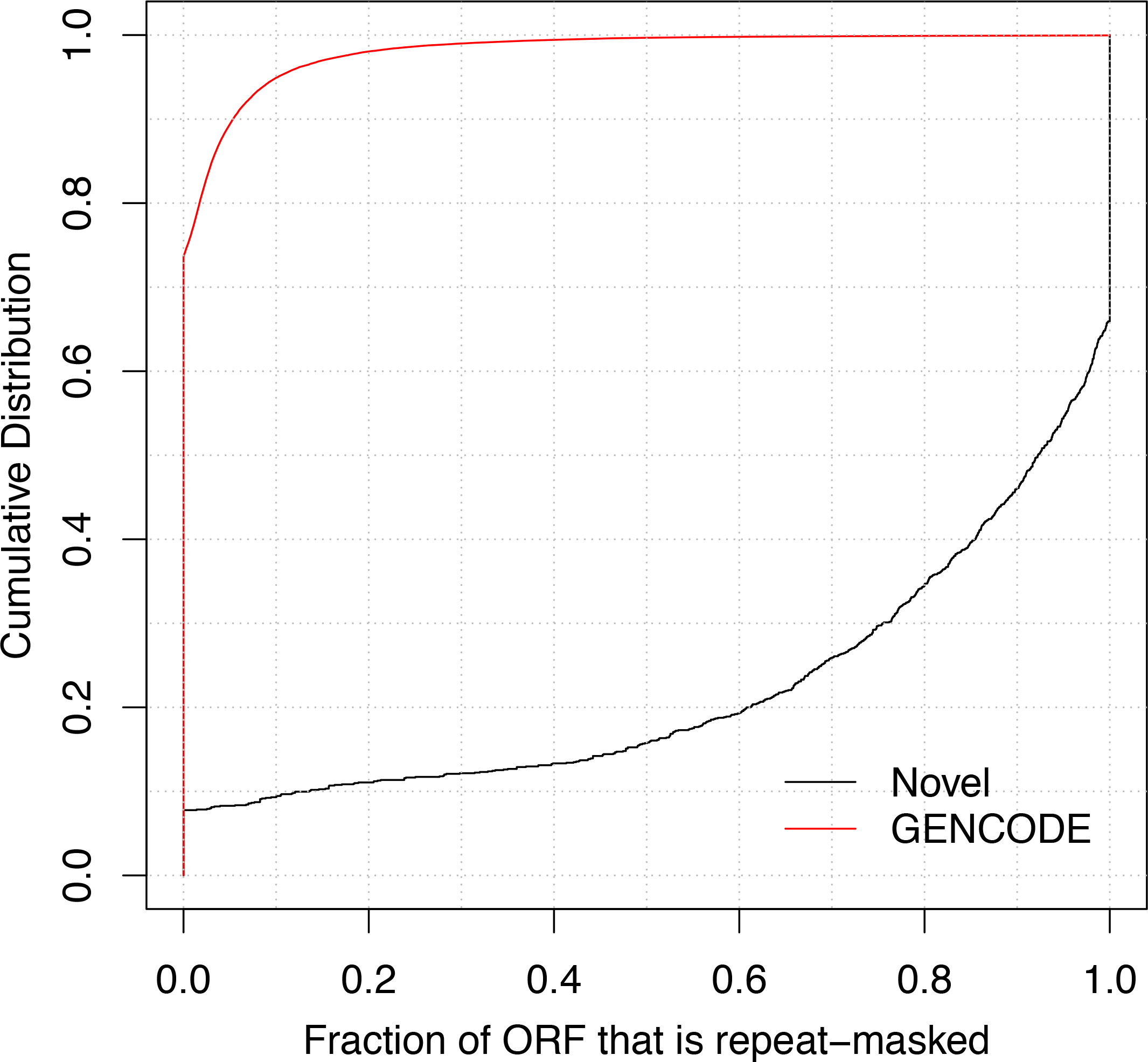
Cumulative distribution of the fraction of each ORF that overlaps RepeatMasker regions for predicted novel coding transcripts (black) and GENCODE protein-coding transcripts (red). The ORFs of over 92% of the predicted novel coding transcripts have some overlap with a RepeatMasker region, including 34% that are entirely contained in RepeatMasker regions, whereas only around 25% of GENCODE coding transcripts overlap repeat regions and in most cases only a small fraction of the transcript overlaps.

**Table 1.**
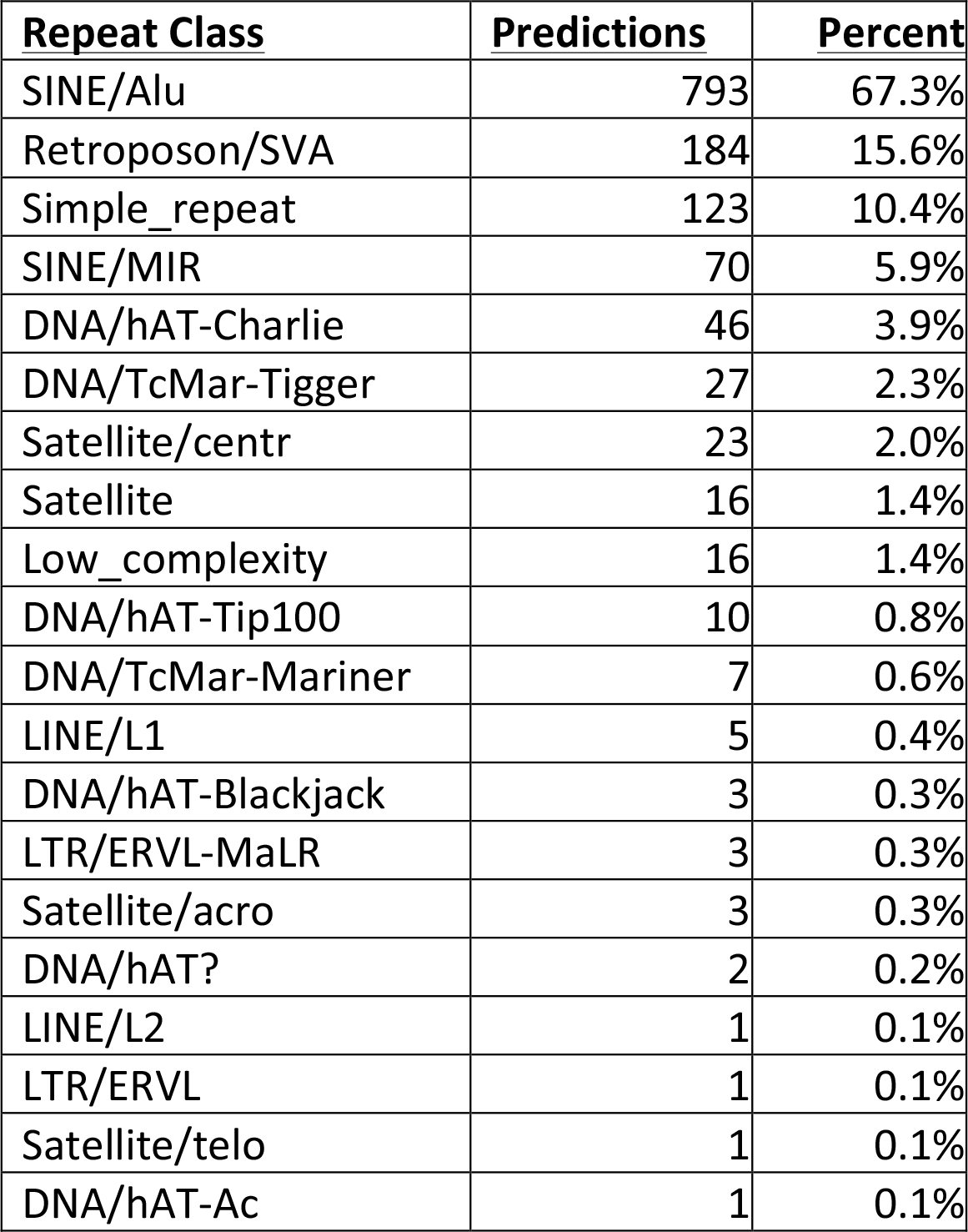
For each repeat class, the number and percent of novel protein-coding predictions that overlap at least one RepeatMasker region of that class. Percentages add up to more than 100% because some loci overlap more than one class.

### Likely pseudogenes

Even if an open reading frame has significant homology to a true protein-coding gene, it could be a disabled copy. To guard against this, GENCODE annotation guidelines consider a paralog to be a pseudogene if there is *any* truncation at all without independent evidence of translation. In the CHESS database, predicted genes are classified as coding if the ORF is at least 75% as long as that of the homology target. However, many pseudogenes would not be excluded by this condition. A nonsense mutation or frame shifting indel in the final 25% of a coding gene would leave more than 75% intact but may break or remove one or more domains at the protein C-terminal, or interfere with proper protein folding. A frame-shifting indel earlier in the sequence could damage an even larger fraction of the protein yet still result in a long ORF by chance. It is common for a disabled paralog to have an ORF more than 75% as long as the parent; in fact, an analysis of a subset of the pseudogenes in the current GENCODE catalog found that 14% of them had ORFs more than 75% as long as those of the parent genes.

Although we believe that most of the novel protein-coding predictions are not homologs of real coding genes, we have found at least two that appear to be disabled paralogs of real coding genes. CHS.625 has aflatoxin B1 aldehyde reductase member 4 homology, as noted by Pertea et al., but the locus contains only the final two coding exons of other family members. According to GENCODE annotation criteria, a duplicated locus with such a large truncation would not be annotated as protein-coding without orthogonal experimental support for its functionality.CHS.56318 is a retrotransposed paralog of *CEBPZOS*. It has a frameshift relative to the parent that replaces the final 10 amino acids with an alternative sequence 21 amino acids long. This changes a substantial fraction of this 80-amino acid protein, whose C-terminal is highly conserved in mammals, so the paralog is unlikely to be coding.

### Lack of protein-coding evolutionary signatures

Next, we tested the CHESS novel ORFs for the evolutionary signature of functional protein-coding regions using PhyloCSF (Lin et al. 2011). PhyloCSF uses substitutions and codon frequencies among 58 mammals to distinguish coding from noncoding regions, with higher scores indicating regions more likely to be translated and functional at the amino acid level. It is often used to distinguish coding transcripts from lncRNAs (Uszczynska-Ratajczak et al. 2018; He et al. 2018). We computed the PhyloCSF score per codon, averaged over the length of the ORF, for the ORFs in all 1,365 predicted novel coding transcripts. As a coding control, we used an equal number of the previously annotated coding ORFs in the CHESS database. As a noncoding control, we used an equal number of theoretical ORFs at least 60 codons long in GENCODE lncRNAs, chosen so as to match the distribution of phylogenetic branch lengths of the species present in the local alignments of the CHESS novel ORFs. This last step was needed because the CHESS novel ORFs tend to have lower alignment branch length than typical lncRNAs, which can decrease the absolute value of the PhyloCSF score. This is notable because usually protein-coding regions have *higher* alignment branch length than noncoding regions, since the latter are so poorly conserved that alignment algorithms can only detect orthology in closely related species. The PhyloCSF score distribution for the CHESS novel ORFs is nearly identical to the distribution for noncoding lncRNAs and much lower than for the previously annotated coding ORFs (Fig. 3), indicating that the novel ORFs are no more likely to be conserved coding regions than lncRNAs currently annotated as noncoding. Even among the 106 novel coding transcripts that do not overlap RepeatMasker regions, most scores are negative, indicating a lack of coding potential.

**Figure 3.**
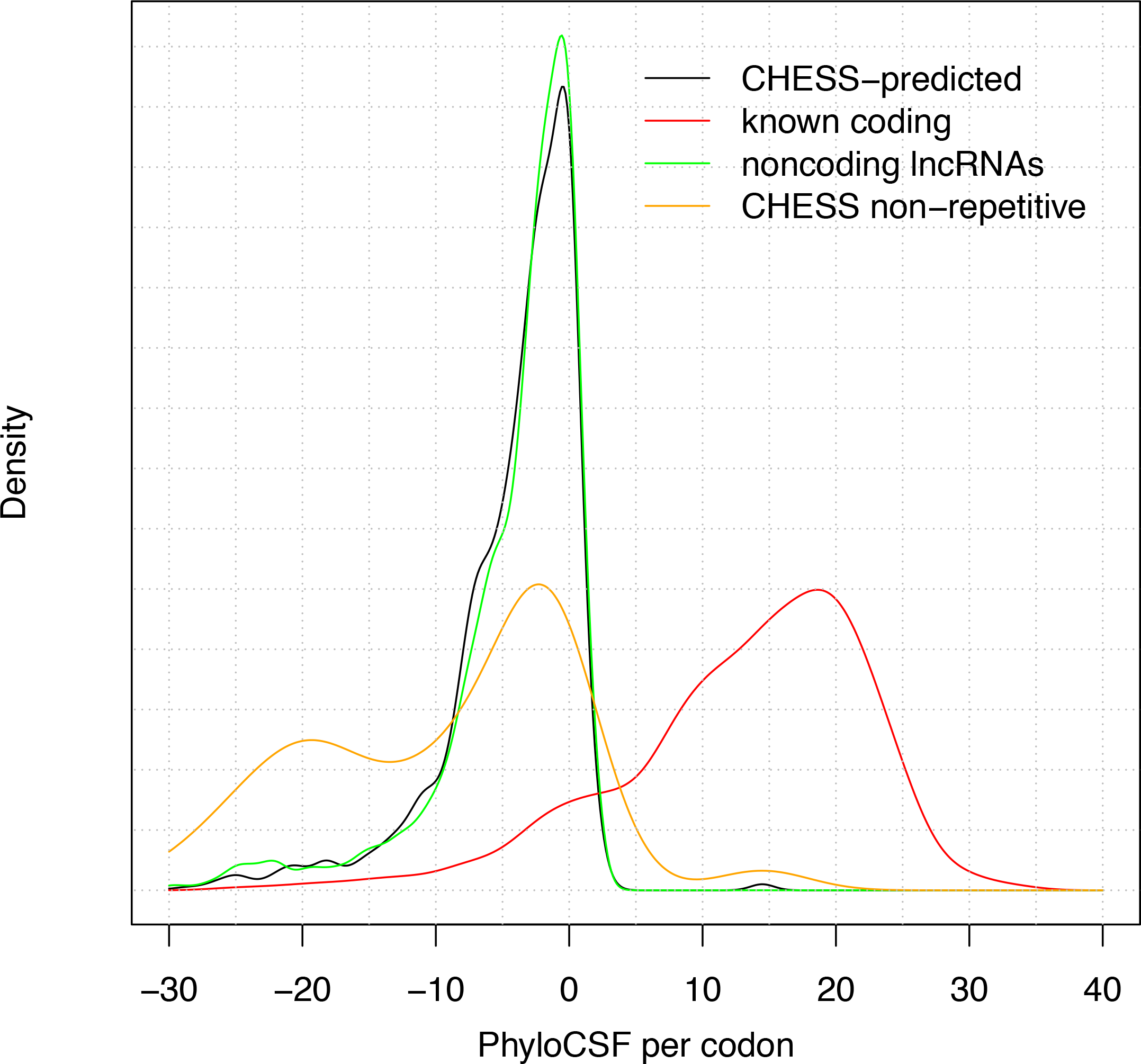
Coding potential of CHESS novel ORFs. Evolutionary coding potential measured by PhyloCSF per codon for novel ORFs (black), previously annotated coding ORFs (red), noncoding theoretical ORFs in Gencode lncRNAs (green), and novel ORFs that do not overlap RepeatMasker regions (orange). The score distribution of the novel ORFs is nearly identical to that of the noncoding lncRNAs, indicating that the novel ORFs are unlikely to be conserved coding regions. Even the novel ORFs that do not overlap RepeatMasker regions lack coding potential.

Only one of the novel protein-coding predictions, CHS.16591, has a high PhyloCSF score. Its ORF is the 3’ end of a longer protein-coding ORF, that of transcript ENST00000497872.4, which was independently added to GENCODE in version 27 (August 2017). This is not a novel gene but is an alternatively spliced transcript of immunoglobulin gene *IGHA2*, which was present in earlier versions of GENCODE.

The lack of a PhyloCSF signal does not itself prove that a gene is noncoding, since roughly 10% of annotated genes do not have such a signal. One explanation for this is that PhyloCSF depends on multi-species genome alignments, and it is therefore not suitable for judging the coding capability of recently-duplicated protein-coding genes (i.e., loci that lack one-to-one orthologs across a reasonable period of evolutionary time). Thus, it is plausible that certain novel coding predictions may score poorly with PhyloCSF because they are lineage-specific paralogs within known gene families as opposed to truly noncoding. However, we doubt that this is a common scenario because our manual analysis has thus far identified just four novel coding predictions that have true homology to genuine protein-coding genes, these being CHS.7402 (the gene presented in Figure 1 of the Pertea manuscript, which GENCODE expects to annotate as a protein-coding gene in a future release), CHS.16591 (which had already been annotated as part of *IGHA2)*, and the two pseudogenes discussed earlier.

### Lack of other evidence of translation

Two additional lines of evidence for protein-coding function were presented in Pertea et al.

First, unmatched spectra from mass spectrometry experiments using 30 human tissues or cell types were searched for potential matches to the novel protein-coding predictions and verified using synthetic spectra. Four matching peptides were identified, each confirming one of the 1,178 novel coding predictions. However, despite the verification procedure these four peptides do not confer a high degree of certainty because each is only seven amino acids long and only one peptide was found per protein. The Human Proteome Project (Deutsch et al. 2016) requires at least two non-overlapping nine-residue peptides to validate a novel coding gene.

In any case, none of these four novel coding predictions has a conserved ORF. Even though CHS.57705 and CHS.24083 have homology to other primate sequences (Pertea et al. 2018), the start codon of CHS.57705 is only preserved in the closest human relatives and CHS.24083 has a premature termination codon in all sequences apart from gorilla (**Fig. 4**). CHS.53541 has a 2-base deletion in apes that changes the reading frame relative to other primates (**Fig. 5a**) and is unlikely to be coding anyway because it has similarity to the GVQW domain proteins and is likely derived from an *Alu* SINE element. Finally, all non-human sequences related to CHS.16287 have a premature termination codon, and several also have frame shifts and missing start codons (**Fig. 5b**).

**Figure 4.**
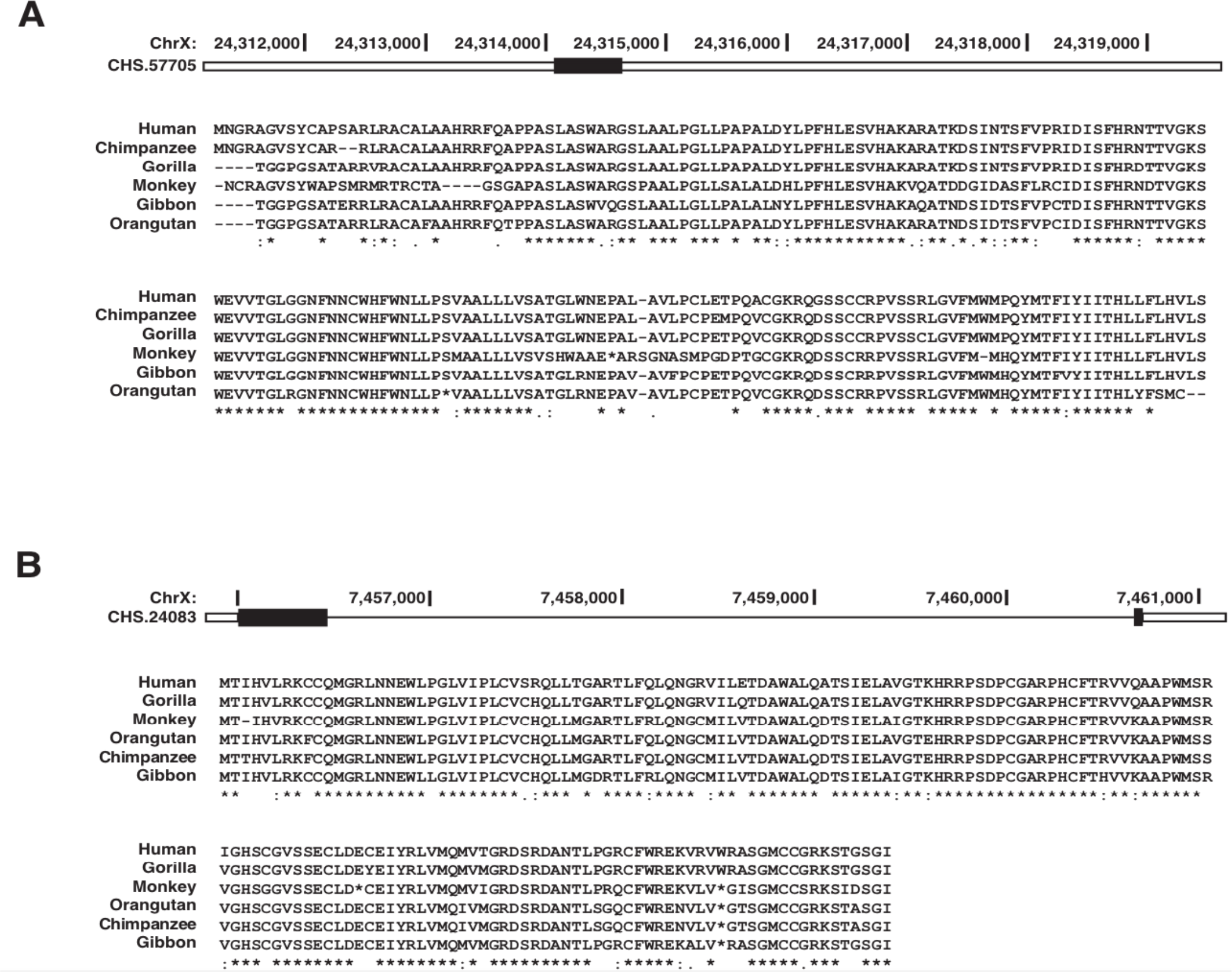
The alignments from Pertea et al. Figure 4 (Pertea et al. 2018) show that the start codon of CHS.57705 is not well conserved and that most species have early stop codons (asterisks) that truncate the final 17 amino acids of CHS.24083. Neither of these open reading frames are conserved in primates.

**Figure 5.**
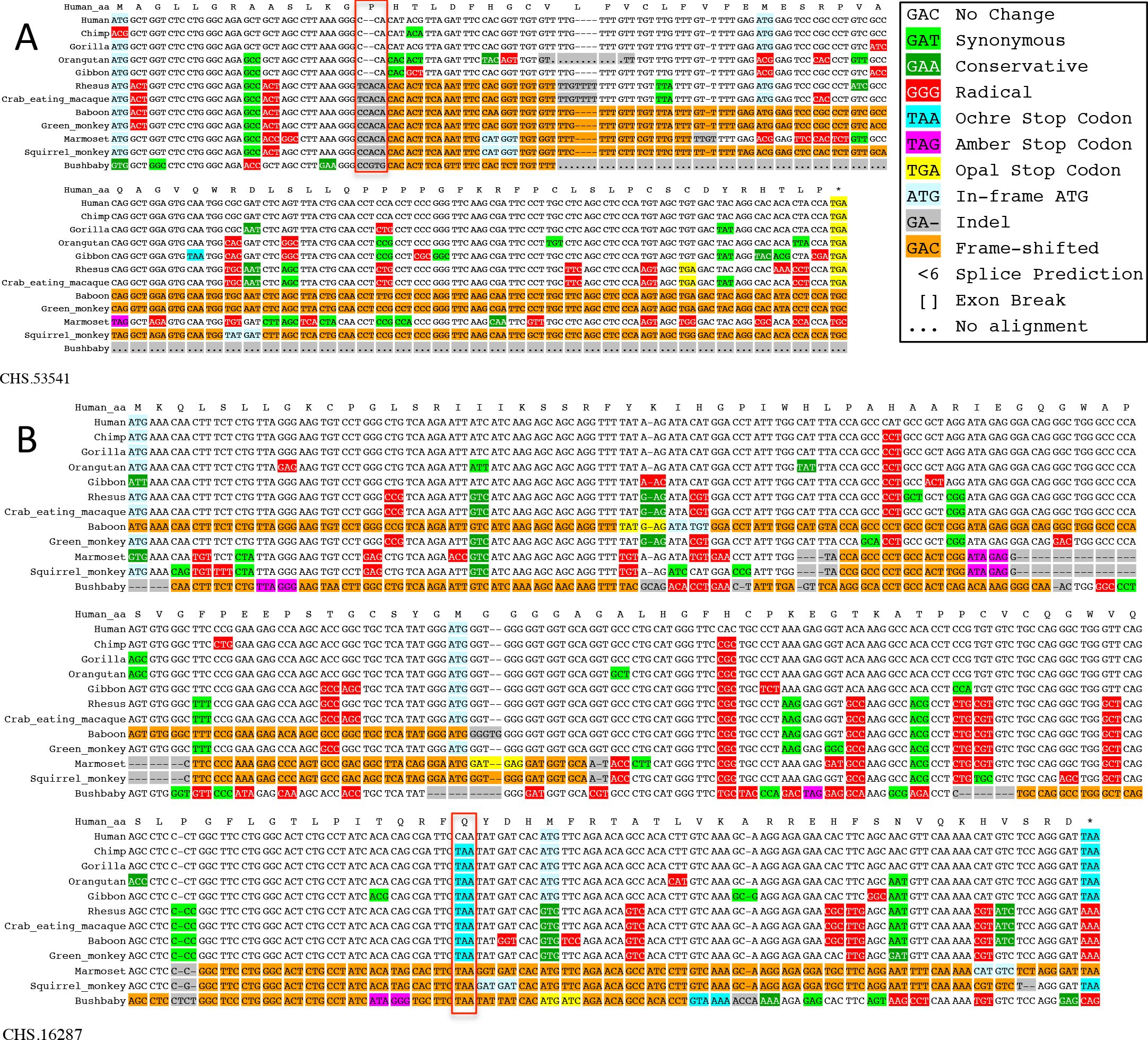
Alignments indicate ORFs are not conserved in primates. (a) A 2-base deletion (red box) in apes in the 14th codon of CHS.57705 changes the reading frame relative to other primates. Also note that the start codon is not present in chimp. (b) A TAA stop codon (red rectangle around cyan codons) truncates the final 29 amino acids of the ORF of CHS.16287. Also note that start codon is not present in gibbon, and much of the ORF is frame-shifted in several species (orange).

We note that Pertea et al. describe CHS.57705 and CHS.24083 as being conserved in primates, whereas we have described them here as not conserved. This discrepancy is due to different usage of the word “conserved”. Pertea et al. use it to mean that the homologous sequences are similar, whereas we are using it to mean that there is evidence of purifying selection. This latter usage requires not only that the sequences be similar but that they are more similar than would be expected under neutral evolution, which in turn requires comparison to a null model of neutral sequence evolution as well as assuring preservation of start codon, stop codon, and reading frame.

Pertea et al. provided evidence of differential expression as a second method to validate the novel protein-coding predictions. However, while differential expression might potentially be an indication that an RNA molecule is functional, it is not an indication that it functions at the protein level. In fact, protein-coding genes tend to be *less* tissue specific than lncRNAs (Cabili et al. 2011).

## Discussion

The integration of massive data sets of transcriptional information with predictions of protein-coding potential is a promising approach to guide the discovery of previously unannotated protein-coding genes, with the understanding that in extensively scrutinized genomes like human finding even a handful of novel protein-coding genes is challenging. In reprocessing a large number of high quality and biologically interesting RNAseq datasets from GTEx, Pertea et al. have created an important starting point for this investigation. However, the subsequent process of protein-coding annotation suffers from a number of flaws that have resulted in a large number of false positives. The discovery of hundreds of novel human protein-coding genes is an extraordinary claim that must be backed up by strong experimental or evolutionary evidence. However the 1,178 CHESS novel protein-coding predictions have little support for coding potential beyond homology to other predictions that are themselves questionable and do not hold up to scrutiny. Our reanalysis of the evidence indicates that nearly all of the novel coding predictions lack the evolutionary signatures of protein-coding genes, stem from homology to transposons, and are unlikely to represent functional protein-coding genes.

### Can transposon-associated sequences code for protein?

Since so many of the novel protein-coding predictions are associated with transposons, it is important to consider whether there is evidence that these can be protein-coding. It is true that a number of human coding genes in GENCODE and RefSeq are derived from transposons (Riordan and Dupuy 2013), but in all cases these genes are well established and have cross-mammalian conservation, whereas the novel coding predictions based on transposons have little to no conservation. Also, almost all of the transposon-derived protein-coding genes that are recognized in human feature coding sequences that have evolved from protein-coding regions contained in the transposon (Riordan and Dupuy 2013). For example, members of the ancient *ZBED* gene family incorporate transposase domains contributed by *hAT* DNA transposons (Hayward et al. 2013). In contrast, more than two thirds of the novel coding predictions overlap *Alu* elements. In common with other members of the SINE family, *Alu* elements do not contain protein-coding regions; instead, the core sequence is thought to be derived from a noncoding signal recognition particle RNA gene. Thus, the only way the transposon-associated novel coding predictions could contain genuine coding sequences is if the majority were *de novo* emergences from noncoding sequences.

We are aware of just one previous study that supports the coding status of primate-derived transposon-based ORFs (Toll-Riera et al. 2009). Toll-Riera et al. identified 154 genes derived from LINE or SINE transposons during primate evolution that were classified as coding in Ensembl. However, the authors used Ensembl version 48, which predated the GENCODE expert annotation process. Today, we observe that over 90% of the transposon-based genes from this paper are no longer classified as coding by GENCODE, while the remaining 13 were all tagged as potential noncoding in a recent study (Abascal et al. 2018). Four of these 13 genes have already been reclassified as noncoding by human experts and the other 9 are under review. Nonetheless, two of these genes - *LINC00269* and *C8orf44* - are still annotated in reference sets and were used to identify 137 novel coding predictions in the CHESS database based on their classification as protein-coding loci.

The authors of the CHESS database did recognize that at least 132 of their novel ORFs overlapped transposons (62% of these were similar to GenBank prediction LP3428), but stated that these novel ORFs could be coding because “many retroposed genes have been previously reported as functional, particularly those that exhibit testis-biased expression”. However, the referenced paper (Emerson et al. 2004) specifically supports the functionality of retrotransposed protein-coding genes. These arise when mRNAs are inserted into the genome sequence through the activity of enzymes encoded by retrotransposons; they are not in themselves transposon-based ORFs. Also, the testis bias reported by Emerson et al only applied to those retrotransposed copies of coding genes that were on the X chromosome, and only 8 of the 132 transposon-based ORFs recognized by Pertea et al. are on the X chromosome.

### Recommendations for future studies

A plethora of recent papers proclaim the discovery of hundreds or thousands of new human protein-coding ORFs (see Introduction), so it seems likely that there will be further studies of this type. For that reason we would like to make several recommendations that authors and reviewers alike might like to bear in mind with the aim of achieving higher confidence protein-coding predictions.

First, data should be filtered for the complete list of transposons. Second, ORFs predicted to be protein-coding based on homology should extend the full length of the coding homolog, unless there is independent evidence of functional translation, to avoid inclusion of pseudogenes. Third, any homology must be to manually-curated genes, not to predicted genes. Fourth, expression at the transcript level is not protein-coding evidence; even ribosome profiling data is not in itself proof of translation into a functional protein. Fifth, be conservative when attributing protein evidence from proteomics experiments; most novel protein coding genes will be hard to detect in standard proteomics experiments because they are likely to be expressed only in low quantities or in limited tissues, but using less stringent thresholds to compensate for that is likely to result in many false positives. Sixth, conservation among related species should be tested against a null model defined by noncoding regions in order to detect purifying selection. Finally it is important that all novel predictions are manually inspected, and not just a select few. Authors should not implicitly trust their own predictions. For example, manual inspection of the CHESS novel protein-coding predictions has quickly revealed that most are based on homology to the same few annotations, many of which are low quality predictions.

Finally, to achieve the level of quality needed for reference gene catalogs, annotation should be supported by the intersection of multiple orthogonal datasets. For example, GENCODE recently added 16 novel human protein-coding genes from mass spectrometry experiments (Wright et al. 2016) and another 139 were added to GENCODE versions 24-28 (December 2015 through April 2018) through a process that combined evolutionary signatures, experimental evidence, and expert manual annotation (Mudge et al. In preparation). In both cases the manual step was essential to weed out false positives.

The GENCODE gene set continues to be updated, and GENCODE does not claim that the current catalog of human protein-coding and noncoding genes is complete. For example, 67, 27, and 43 protein-coding genes were newly annotated and 180, 7, and 41 were deprecated in the last three GENCODE versions (v26-v28, released in March 2017, August 2017, and April 2018, respectively). There are likely to be additional protein-coding genes remaining to be discovered, and certain protein-coding genes currently annotated may be false predictions. The GENCODE Consortium looks forward to continued interactions with the community to incorporate new methods and results with the aim of narrowing the gap between the current reference gene annotation sets and the complex biological reality it aims to represent. As just one example, the set of lncRNAs recently produced by the FANTOM project (Hon et al. 2017) were incorporated into the GENCODE capture long-read sequencing experimental validation pipeline (Lagarde et al. 2017) with the goal of eventually deriving the full length structure of potentially novel lncRNAs yet to be annotated in GENCODE. Similarly, The Consortium is eager to better understand and learn from those who are using GENCODE gene annotation and encourage direct communication via.

We hope that our observations here will lead to improvements in the CHESS methodology and database, and also help guide future large-scale experiments and computational analyses carried out by others.

## Methods

All references to the Pertea et al. manuscript, its supplementary material, and the CHESS database refer to versions downloaded on June 1, 2018. Transcripts were extracted from the GFF file downloaded from http://ccb.jhu.edu/chess/data/chess2.0.gff.gz.

RepeatMasker regions were obtained from the UCSC genome browser at http://hgdownload.soe.ucsc.edu/goldenPath/hg38/bigZips/hg38.fa.out.gz. GENCODE overlaps were calculated using GENCODE version 28 (April 2018). When calculating the fraction of overlap with APPRIS principal isoforms, we used all transcripts tagged as “appris_principal_1”.

PhyloCSF was run using the 58 mammals parameters and the “mle” and “bls” options on the ORF of each transcript, excluding the stop codon, and the resulting score was divided by the length of the ORF in codons to obtain the score per codon. Alignments were extracted from the 100-vertebrate MULTIZ hg38 alignment obtained from the UCSC genome browser (Casper et al. 2017), with species restricted to the 58 placental mammals. Coding controls were a randomly chosen subset of 1365 protein-coding transcripts from the CHESS database that were not in genes with status “novel” or “known_fantom”. To choose noncoding controls, we found the longest theoretical ORF in each lncRNA transcript in GENCODE v28, rejecting any that were not at least 60 codons long. We then chose the subset of 1365 that best matched the phylogenetic branch lengths of the species present in the local alignments of the CHESS novel coding transcripts. Among the CHESS novel coding transcripts there were three whose per-codon PhyloCSF score was approximately 14.5, which is near the middle of the coding distribution, those being three transcripts of CHS.16591. All others had score less than 3, which is within the noncoding distribution.

We estimated the fraction of GENCODE pseudogenes whose longest ORF is more than 75% as long as those of the parents using the subset consisting of the 1300 pseudogenes whose name is the name of a coding gene, followed by the letter P, followed by an integer (e.g., RPL31P2, which is RPL31 pseudogene 2), because these had unambiguous parents.

Alignments in Fig. 5 were color-coded using CodAlignView (I Jungreis, MF Lin, CS Chan, M Kellis 2016).

Manual analysis of genes was carried out according to GENCODE annotation guidelines, documented here: ftp://ftp.sanger.ac.uk/pub/project/havana/Guidelines/Guidelines_March_2016.pdf

Counts of genes added (or deprecated) in various GENCODE versions were determined by counting protein-coding genes in one version for which none of the CDSs of their transcripts overlapped the CDS of any protein-coding transcript in the previous (or next) version.

## Supplementary Material

Supplementary Table 1 is a tab-delimited file which includes the information from the first tab of Pertea et al. supplementary file CHESS_genes.xlsx (which lists all of the CHESS protein-coding predictions with genomic location and description) plus the following columns:

- “Group” indicates group to which novel ORF belongs according to similarity.
- “PerteaTransposon” indicates protein-coding predictions listed as over-expressed in testis and overlapping retroposons in Pertea et al. Supplementary Table S5.
- “RepeatOverlaps” lists all repeat classes reported by RepeatMasker that overlap the protein-coding prediction, in decreasing order of size of overlap.
- “FractionRepeat” has the fraction of all nucleotide positions in ORFs of the protein-coding prediction that are in a RepeatMasker region.
- “FractionTransposon” has the fraction of all nucleotide positions in ORFs of the protein-coding prediction that are in a transposon-associated RepeatMasker region, i.e., one whose repeat class is not Simple_repeat or Low_complexity and does not start with Satellite.

## Acknowledgements

Research reported in this publication was supported by the National Human Genome Research Institute of the National Institutes of Health under Award Number U41HG007234. The content is solely the responsibility of the authors and does not necessarily represent the official views of the National Institutes of Health.

## References

Abascal F, Juan D, Jungreis I, Martinez L, Rigau M, Rodriguez JM, Vazquez J, Tress ML. 2018. Loose ends: almost one in five human genes still have unresolved coding status. Nucleic Acids Res. http://dx.doi.org/10.1093/nar/gky587.

Afink GB, Veenboer G, de Randamie J, Keijser R, Meischl C, Niessen H, Ris-Stalpers C. 2010. Initial characterization of C16orf89, a novel thyroid-specific gene. Thyroid 20: 811–821.

Bazzini AA, Johnstone TG, Christiano R, Mackowiak SD, Obermayer B, Fleming ES, Vejnar CE, Lee MT, Rajewsky N, Walther TC, et al. 2014. Identification of small ORFs in vertebrates using ribosome footprinting and evolutionary conservation. EMBO J 33: 981–993.

Benson DA, Cavanaugh M, Clark K, Karsch-Mizrachi I, Lipman DJ, Ostell J, Sayers EW. 2013. GenBank. Nucleic Acids Res 41: D36–42.

Cabili MN, Trapnell C, Goff L, Koziol M, Tazon-Vega B, Regev A, Rinn JL. 2011. Integrative annotation of human large intergenic noncoding RNAs reveals global properties and specific subclasses. Genes Dev 25: 1915–1927.

Casper J, Zweig AS, Villarreal C, Tyner C, Speir ML, Rosenbloom KR, Raney BJ, Lee CM, Lee BT, Karolchik D, et al. 2017. The UCSC Genome Browser database: 2018 update. Nucleic Acids Res. http://dx.doi.org/10.1093/nar/gkx1020.

Crappé J, Van Criekinge W, Trooskens G, Hayakawa E, Luyten W, Baggerman G, Menschaert G. 2013. Combining in silico prediction and ribosome profiling in a genome-wide search for novel putatively coding sORFs. BMC Genomics 14: 648.

Deutsch EW, Overall CM, Van Eyk JE, Baker MS, Paik Y-K, Weintraub ST, Lane L, Martens L, Vandenbrouck Y, Kusebauch U, et al. 2016. Human Proteome Project Mass Spectrometry Data Interpretation Guidelines 2.1. J Proteome Res 15: 3961–3970.

Emerson JJ, Kaessmann H, Betrán E, Long M. 2004. Extensive gene traffic on the mammalian X chromosome. Science 303: 537–540.

Ezkurdia I, Calvo E, Del Pozo A, Vázquez J, Valencia A, Tress ML. 2015. The potential clinical impact of the release of two drafts of the human proteome. Expert Rev Proteomics 12: 579–593.

Finn RD, Bateman A, Clements J, Coggill P, Eberhardt RY, Eddy SR, Heger A, Hetherington K, Holm L, Mistry J, et al. 2014. Pfam: the protein families database. Nucleic Acids Res 42: D222–30.

Gascoigne DK, Cheetham SW, Cattenoz PB, Clark MB, Amaral PP, Taft RJ, Wilhelm D, Dinger ME, Mattick JS. 2012. Pinstripe: a suite of programs for integrating transcriptomic and proteomic datasets identifies novel proteins and improves differentiation of protein-coding and non-coding genes. Bioinformatics 28: 3042–3050.

Guttman M, Russell P, Ingolia NT, Weissman JS, Lander ES. 2013. Ribosome profiling provides evidence that large noncoding RNAs do not encode proteins. Cell 154: 240–251.

Hancks DC, Kazazian HH Jr. 2010. SVA retrotransposons: Evolution and genetic instability. Semin Cancer Biol 20: 234–245.

Harrow J, Frankish A, Gonzalez JM, Tapanari E, Diekhans M, Kokocinski F, Aken BL, Barrell D, Zadissa A, Searle S, et al. 2012. GENCODE: the reference human genome annotation for The ENCODE Project. Genome Res 22: 1760–1774.

Hayward A, Ghazal A, Andersson G, Andersson L, Jern P. 2013. ZBED evolution: repeated utilization of DNA transposons as regulators of diverse host functions. PLoS One 8: e59940.

He Q, Liu Y, Sun W. 2018. Statistical analysis of non-coding RNA data. Cancer Lett 417: 161–167.

Hon C-C, Ramilowski JA, Harshbarger J, Bertin N, Rackham OJL, Gough J, Denisenko E, Schmeier S, Poulsen TM, Severin J, et al. 2017. An atlas of human long non-coding RNAs with accurate 5’ ends. Nature 543: 199–204.

I Jungreis, MF Lin, CS Chan, M Kellis. 2016. CodAlignView. CodAlignView: The Codon Alignment Viewer. http://data.broadinstitute.org/compbio1/cav.php (Accessed April 30, 2016).

Kim M-S, Pinto SM, Getnet D, Nirujogi RS, Manda SS, Chaerkady R, Madugundu AK, Kelkar DS, Isserlin R, Jain S, et al. 2014. A draft map of the human proteome. Nature 509: 575–581.

Lagarde J, Uszczynska-Ratajczak B, Carbonell S, Pérez-Lluch S, Abad A, Davis C, Gingeras TR, Frankish A, Harrow J, Guigo R, et al. 2017. High-throughput annotation of full-length long noncoding RNAs with capture long-read sequencing. Nat Genet 49: 1731–1740.

Lassmann T, Frings O, Sonnhammer ELL. 2009. Kalign2: high-performance multiple alignment of protein and nucleotide sequences allowing external features. Nucleic Acids Res 37: 858–865.

Lin MF, Jungreis I, Kellis M. 2011. PhyloCSF: a comparative genomics method to distinguish protein coding and non-coding regions. Bioinformatics 27: i275–82.

MacArthur DG, Balasubramanian S, Frankish A, Huang N, Morris J, Walter K, Jostins L, Habegger L, Pickrell JK, Montgomery SB, et al. 2012. A systematic survey of loss-of-function variants in human protein-coding genes. Science 335: 823–828.

Mackowiak SD, Zauber H, Bielow C, Thiel D, Kutz K, Calviello L, Mastrobuoni G, Rajewsky N, Kempa S, Selbach M, et al. 2015. Extensive identification and analysis of conserved small ORFs in animals. Genome Biol 16: 179.

Marchler-Bauer A, Bo Y, Han L, He J, Lanczycki CJ, Lu S, Chitsaz F, Derbyshire MK, Geer RC, Gonzales NR, et al. 2017. CDD/SPARCLE: functional classification of proteins via subfamily domain architectures. Nucleic Acids Res 45: D200–D203.

Mudge JM, Jungreis I, Hunt T, Gonzalez JM, Wright J, Kay M, Davidson C, Fitzgerald S, Seal R, Tweedie S, et al. In preparation. A new workflow built on whole-genome PhyloCSF finds 144 high-confidence novel conserved protein-coding genes, with many disease associations.

Nesvizhskii AI. 2014. Proteogenomics: concepts, applications and computational strategies. Nat Methods 11: 1114–1125.

O’Leary NA, Wright MW, Brister JR, Ciufo S, Haddad D, McVeigh R, Rajput B, Robbertse B, Smith-White B, Ako-Adjei D, et al. 2016. Reference sequence (RefSeq) database at NCBI: current status, taxonomic expansion, and functional annotation. Nucleic Acids Res 44: D733–45.

Pertea M, Shumate A, Pertea G, Varabyou A, Chang Y-C, Madugundu AK, Pandey A, Salzberg S. 2018. Thousands of large-scale RNA sequencing experiments yield a comprehensive new human gene list and reveal extensive transcriptional noise. bioRxiv 332825. https://www.biorxiv.org/content/early/2018/05/29/332825 (Accessed June 2, 2018).

Riordan JD, Dupuy AJ. 2013. Domesticated transposable element gene products in human cancer. Mob Genet Elements 3: e26693.

Rodriguez JM, Rodriguez-Rivas J, Di Domenico T, Vázquez J, Valencia A, Tress ML. 2018. APPRIS 2017: principal isoforms for multiple gene sets. Nucleic Acids Res 46: D213–D217.

Slavoff SA, Mitchell AJ, Schwaid AG, Cabili MN, Ma J, Levin JZ, Karger AD, Budnik BA, Rinn JL, Saghatelian A. 2013. Peptidomic discovery of short open reading frame-encoded peptides in human cells. Nat Chem Biol 9: 59–64.

Smit AFA, Hubley R, Green P. 2013. 2013–2015. RepeatMasker Open-4.0.

The GTEx Consortium. 2015. The Genotype-Tissue Expression (GTEx) pilot analysis: Multitissue gene regulation in humans. Science 348: 648–660.

The UniProt Consortium. 2017. UniProt: the universal protein knowledgebase. Nucleic Acids Res 45: D158–D169.

Toll-Riera M, Bosch N, Bellora N, Castelo R, Armengol L, Estivill X, Albà MM. 2009. Origin of primate orphan genes: a comparative genomics approach. Mol Biol Evol 26: 603–612.

Uszczynska-Ratajczak B, Lagarde J, Frankish A, Guigó R, Johnson R. 2018. Towards a complete map of the human long non-coding RNA transcriptome. Nat Rev Genet. http://dx.doi.org/10.1038/s41576-018-0017-y.

Vanderperre B, Lucier J-F, Bissonnette C, Motard J, Tremblay G, Vanderperre S, Wisztorski M, Salzet M, Boisvert F-M, Roucou X. 2013. Direct detection of alternative open reading frames translation products in human significantly expands the proteome. PLoS One 8: e70698.

Verheggen K, Volders P-J, Mestdagh P, Menschaert G, Van Damme P, Gevaert K, Martens L, Vandesompele J. 2017. Noncoding after All: Biases in Proteomics Data Do Not Explain Observed Absence of lncRNA Translation Products. J Proteome Res 16: 2508–2515.

Wang H, Xing J, Grover D, Hedges DJ, Han K, Walker JA, Batzer MA. 2005. SVA elements: a hominid-specific retroposon family. J Mol Biol 354: 994–1007.

Wilhelm M, Schlegl J, Hahne H, Gholami AM, Lieberenz M, Savitski MM, Ziegler E, Butzmann L, Gessulat S, Marx H, et al. 2014. Mass-spectrometry-based draft of the human proteome. Nature 509: 582–587.

Wright JC, Mudge J, Weisser H, Barzine MP, Gonzalez JM, Brazma A, Choudhary JS, Harrow J. 2016. Improving GENCODE reference gene annotation using a high-stringency proteogenomics workflow. Nat Commun 7: 11778.

Zerbino DR, Achuthan P, Akanni W, Amode MR, Barrell D, Bhai J, Billis K, Cummins C, Gall A, Girón CG, et al. 2018. Ensembl 2018. Nucleic Acids Res 46: D754–D761.

